# MSModDetector: A Tool for Detecting Mass Shifts and Post-Translational Modifications in Individual Ion Mass Spectrometry Data

**DOI:** 10.1101/2023.06.06.543961

**Authors:** Marjan Faizi, Ryan T Fellers, Dan Lu, Bryon S Drown, Ashwini Jambhekar, Galit Lahav, Neil L Kelleher, Jeremy Gunawardena

## Abstract

**Motivation:** Post-translational modifications (PTMs) on proteins regulate protein structures and functions. A single protein molecule can possess multiple modification sites that can accommodate various PTM types, leading to a variety of different patterns, or combinations of PTMs, on that protein. Different PTM patterns can give rise to distinct biological functions. To facilitate the study of multiple PTMs, top-down mass spectrometry (MS) has proven to be a useful tool to measure the mass of intact proteins, thereby enabling even widely separated PTMs to be assigned to the same protein molecule and allowing determination of how many PTMs are attached to a single protein.

**Results:** We developed a Python module called MSModDetector that studies PTM patterns from individual ion mass spectrometry (I MS) data. I MS is an intact protein mass spectrometry approach that generates true mass spectra without the need to infer charge states. The algorithm first detects and quantifies mass shifts for a protein of interest and subsequently infers potential PTM patterns using linear programming. The algorithm is evaluated on simulated I MS data and experimental I MS data for the tumor suppressor protein p53. We show that MSModDetector is a useful tool for comparing a protein’s PTM pattern landscape across different conditions. An improved analysis of PTM patterns will enable a deeper understanding of PTM-regulated cellular processes.

**Availability:** The source code is available at https://github.com/marjanfaizi/MSModDetector together with the scripts used for analyses and to generate the figures presented in this study.

## 1 Introduction

Post-translational modifications (PTMs) are covalent modifications of proteins that can affect their activity, stability, and interaction with other components, dependent on both the type and the location of modification on the protein. Proteins can have multiple modification sites with various PTMs that act in combination to influence protein function (Prabakaran et al., 2012; Leutert et al., 2021). We refer to the specific combination of PTMs on a protein molecule as its “modform” (Prabakaran et al., 2012) (Fig. 1A). Proteins harboring multiple modification sites are preferentially found to act as hubs in protein-protein interaction networks and are significantly more associated with human disease than proteins with no known modification sites (Huang et al., 2014). Additionally, PTMs also play an important role in cellular information processing, with protein modforms being able to encode information about upstream conditions which in turn guides downstream responses (Prabakaran et al., 2012). Hence, the successful measurement of co-occurring PTMs, rather than PTMs on single sites, is important to connect more precisely protein function with cellular physiology (Csizmok and Forman-Kay, 2018).

**Figure 1:**
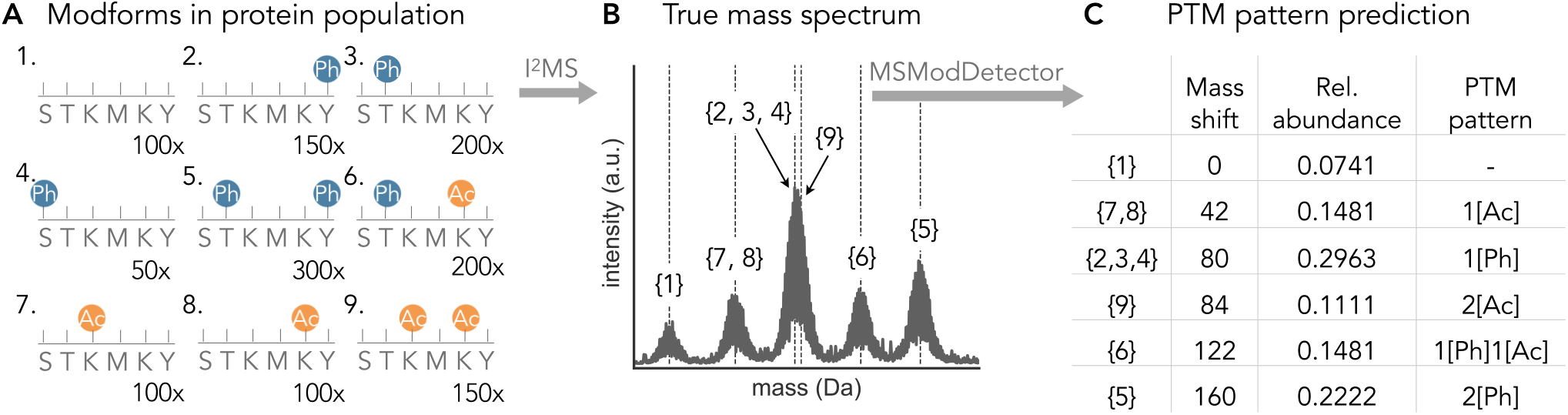
PTM pattern inference problem. The protein population in (A) can be modified by phosphorylation (Ph) at the serine (S), threonine (T), or tyrosine (Y) sites or it can be acetylated (Ac) at the lysine (K) sites, leading to different combinations of PTMs (“modforms”) within the population of the same protein. All 32 possible modforms are listed in Supplementary Fig. S1. The numbers on the bottom right indicate the amount of the respective modform. (B) I MS measures the total mass of each protein molecule and generates a true mass spectrum. Modforms 2, 3, and 4 have the same mass shift and the same isotopic distribution, as do modforms 7 and 8. These “patterns”, which have the same numbers of different modifications, are what is resolved by I MS. The patterns with one phosphorylation (modforms 2, 3, 4) or 2 acetylations (modform 9) can not be resolved visually in the spectrum as their isotopic distributions overlap too closely. (C) The goal of MSModDetector is to identify mass shifts caused by the modifications and to infer which PTM patterns are present and estimate their abundance using linear programming.

Mass spectrometry (MS) has proven to be a powerful tool to determine patterns of PTMs (Jensen, 2006). The development of MS methods has paved the way to study protein modforms (Prabakaran et al., 2012; Leutert et al., 2021), and currently there are two different MS approaches to tackle this problem. The predominant “bottom-up” MS strategy uses proteases to cleave proteins into smaller peptides before mass determination and PTM identification (Aebersold et al, 2018). Identified peptides are then used to infer the presence and modification state of protein groups using tools like MaxQuant (Tyanova et al., 2016) or MSFragger (Kong et al., 2017). This approach is well established and provides high mass accuracy and resolution. However, bottom-up MS is limited in its ability to detect widely separated modifications on the same protein molecule (Schaffer et al., 2019). Due to pre-enzymatic digestion it is impossible to distinguish whether different modification sites on separate peptides originate from a single protein molecule or from distinct molecules (Compton et al., 2018).

An alternative MS approach, called “top-down” MS, aims to overcome this issue by characterizing intact proteins, revealing co-occurring PTMs that may be well separated on an individual protein molecule (Aebersold et al, 2018; Compton et al., 2018). Measuring intact proteins with top-down MS approaches gives rise to complex isotopic distributions in the observed spectrum, expressed in units of mass-to-charge ratio (*m*/*z*). The isotopic distribution is a result of varying amounts of neutrons for the same element. As the mass and number of atoms of a protein increases, the likelihood of observing more heavy isotopes of that protein species increases as well (Schaffer et al., 2019). The electrospray ionization process produces a distribution of highly-charged precursor ions with *m*/*z* values within the detection range of the mass analyzer. The first step to analyse these complex *m*/*z* spectra involves deconvolving the isotopic distribution and resolving the charge states of the peaks to convert the *m*/*z* value into mass values. There are many deconvolution algorithms, e.g., THRASH (Horn et al., 2000) or FLASHDeconv (Jeong et al., 2020), however, this step is often accompanied by a considerable number of missassigned peaks (Cai et al., 2016). A new top-down technology called individual ion mass spectrometry (I^2^MS) has recently been developed (Kafader et al., 2020), which can create a true mass spectrum by measuring the charge directly to determine the mass of each ion. Thus, there is no need to infer the charge state in order to convert the *m*/*z* values into mass values. On its own, without fragmentation of the protein and analysis of the fragments (tandem mass spectrometry), I^2^MS is unable to resolve modforms with the same numbers of modifications, as these have identical mass shifts (Fig. 1B). We refer to such modforms with identical mass shifts as PTM “patterns”; modforms 2, 3 and 4 in Fig. 1A form such a pattern, with just a single phosphorylation. It may further happen that distinct patterns have mass shifts that are sufficiently similar that they cannot be resolved by I^2^MS alone (Fig. 1B).

Commonly available tools for top-down MS data, like MASH Suite Pro (Cai et al., 2016) or TopPIC (Kou et al., 2016), identify and quantify proteins and PTMs based on *m*/*z* spectra and are therefore not suited to process the true mass spectra generated by I^2^MS. Furthermore, identification of unmodified or modified proteins is based on matching the observed spectra to entries in a protein sequence database (Schaffer et al., 2019). If the modified protein is not listed in the database it remains undetected. To overcome this issue, some tools allow a mass offset during the database search and matching process to account for unknown mass shifts (Liu et al., 2012). However, the number of PTMs allowed to explain the unknown mass shifts is usually limited to a small number, e.g., one or two PTMs in the case of TopPIC (Kou et al., 2016). Accordingly, the modforms and PTM patterns of important hub proteins harboring many modification sites cannot be studied with commonly available tools.

Therefore, we developed a Python module, called MSModDetector, to facilitate the analysis of I^2^MS mass spectra and interpretation of PTM patterns. MSModDetector consists of an algorithm that detects mass shifts within the mass spectrum and a linear program to estimate the PTM patterns corresponding to a given mass shift. The algorithm is not limited in how many different PTMs can be used to explain a given mass shift (Fig. 1C). We applied the algorithm to experimental I^2^MS data, obtained from the tumor suppressor protein p53 in MCF7 cells; p53 is a hub protein and known to be modified at many sites (Hafner et al., 2019). We present a table of mass shifts for p53 under different cellular conditions with the corresponding PTM patterns (Table 1).

**Table 1:** PTM pattern predictions for the I MS data of endogenous p53 under Nutlin-3a (10 *μM*) and UV (10 *J/m*) conditions. For each mass shift the relative abundances for both replicates are shown as well as PTM pattern predictions for two different objective functions: “min both” is minimizing the number of PTMs and the error between observed and inferred mass shift and “min ptm” is only minimizing the number of PTMs. Mass shifts detected in more than one condition or replicate are combined by calculating their average (see section 2.1.5). The following PTM types are considered for the pattern combinations: phosphorylation (Ph), acetylation (Ac), methylation (Me1), di-methylation (Me2), tri-methylation (Me3). In addition to biological functional modifications we also consider artifacts like: phosphate (Ph-OH), oxidation (Ox), cysteinylation (Cys), sodium adduct (Na).

| mass shift | PTM pattern |  | rel. abundance |  |  |  |
| --- | --- | --- | --- | --- | --- | --- |
|  | PTM pattern (min both) | PTM pattern (min ptm) | Nutlin-3a (rep.1) | Nutlin-3a (rep. 2) | UV (rep. 1) | UV (rep. 2) |
| 120.7 | 1[Ph-OH]1[Na] | 1[Cys] | 0.13560 | 0.1740 | 0.0864 | 0.0898 |
| 136.2 | 1[Ph]2[Me2] | 1[Cys]1[Ox] | 0.0239 | - | - | - |
| 180.3 | 1[Ph]2[Me3]1[Ox] | 1[Ph]2[Me3]1[Ox] | - | 0.0714 | - | 0.0273 |
| 200.7 | 1[Ph-OH]1[Ph]1[Na] | 1[Cys]1[Ph] | 0.0470 | 0.0730 | 0.0290 | 0.1002 |
| 216.7 | 1[Cys]1[Ph-OH] | 1[Cys]1[Ph-OH] | - | 0.0226 | - | 0.0329 |
| 259.3 | 1[Cys]1[Ph-OH]1[Me3] | 1[Cys]1[Ph-OH]1[Me3] | - | - | - | 0.0366 |
| 297.8 | 2[Ph-OH]1[Ph]1[Na] | 1[Cys]1[Ph-OH]1[Ph] | 0.1578 | 0.1292 | 0.1045 | 0.0710 |
| 309.9 | 3[Ph-OH]1[Ox] | 3[Ph-OH]1[Ox] | - | 0.0265 | - | - |
| 357.4 | 3[Cys] | 3[Cys] | 0.0617 | 0.0763 | 0.0415 | 0.0380 |
| 377.3 | 1[Cys]2[Ph]2[Me3]1[Me1] | 1[Cys]1[Ph-OH]2[Ph] | 0.1045 | 0.1390 | 0.1510 | 0.1359 |

In order to set these results in context and understand how many predicted PTM patterns are correct, we constructed a simulated data set and evaluated the algorithm on it. We introduced noise and error to the simulated data and constructed overlapping isotopic distributions to see how much noise/error or overlap can be tolerated by the algorithm in order to make reasonable predictions. We observe that adding noise or error to the data does not impact the algorithm’s prediction, however overlapping isotopic distributions interfere with the results.

## 2 Materials and methods

The Python module MSModDetector provides an algorithm that takes I^2^MS data as input and determines mass shifts within a pre-selected mass range for a protein of interest. The relative amount of each mass shift is calculated and a PTM pattern is inferred for every detected mass shift using a linear program. For the algorithm to run successfully, the minimum average mass of the protein of interest is required to be 14 kDa (see section 2.1.3 for further details). The algorithm is tested on simulated and experimental I^2^MS data. Both data sources will be described in section 2.3 and 2.4, respectively. Fig. 2 depicts the workflow of the algorithm and a detailed description of the processes of mass shift detection and PTM pattern inference is given in the following sections.

**Figure 2:**
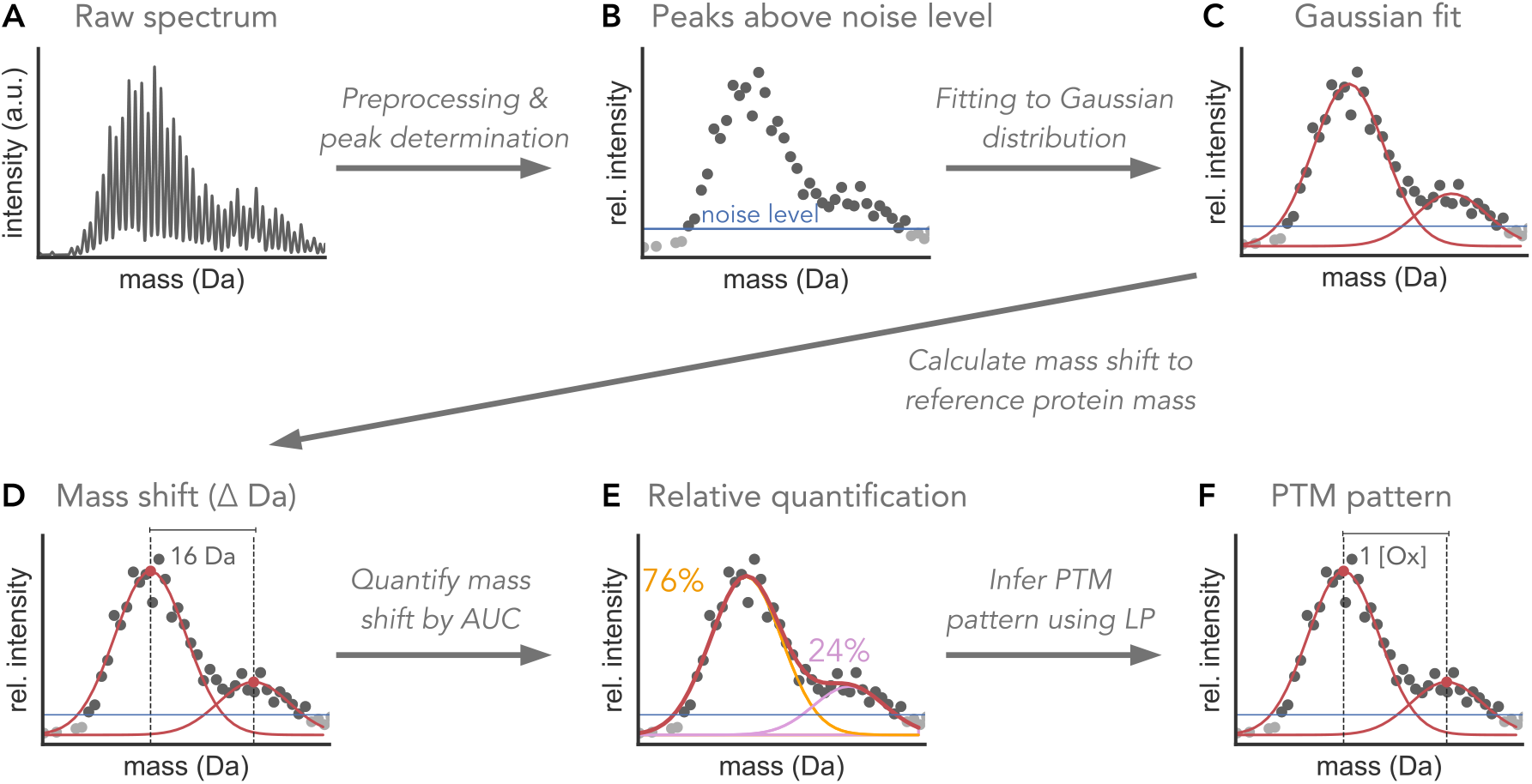
MSModDetector workflow depicted on a toy example. (A) I MS raw data is preprocessed and (B) only peaks above a determined noise level are selected for further analysis. (C) Gaussian distributions are then fitted to the data by sliding a window of fixed size over the spectrum. A chi-square score assesses the goodness of fit and selects for the best fit. (D) The means of the Gaussian distributions are used to calculate the mass shift to the reference protein mass. (E) The area under the curve (AUC) is calculated for each distribution and used to relatively quantify every mass shift in relation to all detected mass shifts. (F) A linear program (LP) infers potential PTM patterns for the given mass shifts. Here, a PTM pattern of one oxygen (Ox) is inferred for a mass shift of 16 Da.

### 2.1 Detection of mass shifts

#### 2.1.1 Raw data preprocessing

Raw I^2^MS data is processed to create mass spectra as described (Melani et al., 2022). The spectra are exported as profile .mzml files and contain mass values in units of Dalton (Da) and their corresponding intensity values which indicate the relative abundances (Fig. 2A). The algorithm uses the profile .mzml files as input and requires a predefined mass range in which the algorithm searches for mass shifts. The mass range should contain the mass of the protein of interest. Intensities of the masses within this range are normalized by the maximum intensity in this window. Normalized intensities facilitate subsequent analyzes and are re-scaled after the analysis. The profile data are further processed by centroiding, which involves retaining only the maximum intensity value of each peak in the mass spectrum. Peaks above a pre-determined noise level are defined as signals and used to determine mass shifts (Fig. 2B). The noise level defined in this study is equal to one-half of the standard deviation of all intensities within the predefined mass range. We found this metric to be a good measure to distinguish signals from background noise.

#### 2.1.2 Gaussian model and isotopic distribution

After preprocessing the raw data, the isotopic distributions in the mass spectrum are fitted by Gaussian functions (Fig. 2C). For this we assume that the isotopic distribution follows a Gaussian distribution and the standard deviation *σ* of the isotopic distribution for a protein of interest is constant and does not change when modifications are considered. The standard deviation *σ* is calculated by fitting a Gaussian function to the isotopic distribution of the unmodified protein of interest. The isotopic distribution of the unmodified protein is calculated with the Python package *pyopenms* and only requires the protein sequence in FASTA format.

The algorithm detects mass shifts by sliding a window of fixed size over the mass spectrum with a step size of 1 Da and fitting Gaussian distributions to the data within this window. Since the standard deviation *σ* is constant only the mean and amplitude are fitted to the data. The user can determine how much overlap between two fitted Gaussian distributions is allowed. We determine the allowed overlap by setting a minimal distance between the mean values of two fitted Gaussian distributions. We recommend the minimum distance to be at least two-thirds of the window size.

#### 2.1.3 Chi-square goodness of fit

The chi-square goodness of fit test is used to evaluate the fit of the Gaussian function to the data. The null hypothesis of the chi-square goodness of fit test assumes that the observed (experimentally-obtained) distribution is the same as the expected (Gaussian fitted) distribution. Therefore, if the p-value is small (less than the pre-determined significance level) we can reject the null hypothesis that both distributions are the same, whereas high p-values reflect a good fit. Hence, for every fit, a chi-square score is calculated and a p-value is assigned after each step. All fitting results with a p-value below the user-defined significance level are removed. Furthermore, if the mean values of two adjacent Gaussian distributions are closer than the user-defined allowed overlap, the fit with the lower p-value is removed as well.

To perform the chi-square test, at least 5 peaks are required within the sliding window. Hence, the size of the sliding window should be bigger than 5 Da to make sure that at least 5 peaks are used to calculate the chi-square score and p-value. We further recommend selecting a window size that covers approximately the upper third part of the isotopic distribution (Supplementary Fig. S2). For proteins with an average mass of at least approximately 14 kDa, we can make sure that the upper third part of the isotopic distribution contains at least 5 peaks (Supplementary Fig. S2B).

#### 2.1.4 Mass shift identification and relative quantification

After the mean values are determined, they are used to calculate the mass shift from the reference mass of the unmodified protein of interest (Fig. 2D). Each mass shift is relatively quantified by calculating the area under the curve of its Gaussian distribution (Fig. 2E). The abundance for a mass shift (*A*) is calculated as the integral of the Gaussian function, 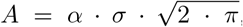, with *α* being the amplitude of the fitted Gaussian function. This approach provides a notion of the abundance of the given mass shift relative to all mass shifts observed within the predefined mass range.

#### 2.1.5 Mass shift alignment across samples

To compare mass shifts across multiple samples, the fitted mean values of all samples are aligned with each other. If for instance the difference in mass for the mean values in two different samples is less than 0.5 Da, then both mean values are assumed to represent the same species and are binned together. The binning size depends on the pre-defined mass tolerance, the maximal accepted error between the observed and inferred mass shift, and can be increased by setting a higher mass tolerance. However, we emphasize that increasing the mass tolerance and considering a larger bin size results in more potential PTM pattern combinations. The mean values are binned and an average mass from all mean values within one bin is calculated. The average mass is then used for mass shift identification.

### 2.2 Inference of PTM patterns

For each mass shift, a linear program (LP) is solved to identify potential combinations of PTM patterns (Fig. 2F). One mass shift can give rise to multiple solutions of PTM patterns. To select for one PTM pattern, the algorithm provides three different objective functions that result in the following LPs: (i) an LP with the objective to minimize the number of total PTMs on a single protein, (ii) an LP with the objective to minimize the error between observed and inferred mass shifts, and (iii) an LP that combines both previously mentioned objective functions. While the objective functions are not biologically motivated, they are still useful and can be seen as a method to rank the multiple solutions or potential PTM patterns. In order to obtain the best *k* optimal solutions, the variable *laps* can be specified by the user which determines the number of times the optimization problem is solved. We prevent obtaining the same solution *k* times by using the optimal solution of the previous run as a constraint for the current run.

#### 2.2.1 Linear program to minimize the number of PTMs

The first LP is based on the objective of minimizing the total number of PTMs, *P*, on a single protein:

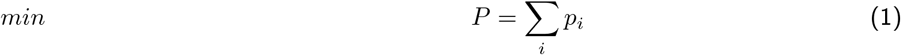

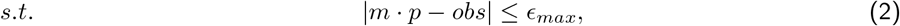

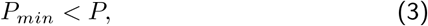

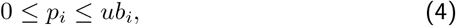

where *m* is a row vector that contains the masses of the different PTMs and *p* is a column vector of the unknown number of each PTM type. Multiplying both vectors results in the mass shift for the inferred PTM pattern. The difference between the theoretical and observed mass shift *obs* is constrained by *ϵ*_*max*_, the mass tolerance defined by the user. Each number of PTMs *p*_*i*_ is a positive number and is constrained by an upper bound *ub*_*i*_. *P*_*min*_ is the optimal solution from the previous result and helps to find the next optimal solution.

#### 2.2.2 Linear program to minimize the mass error

For the second LP we minimize the difference, *ϵ*, between inferred and observed mass shift.

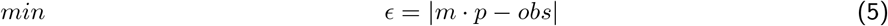

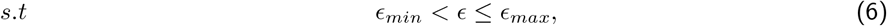

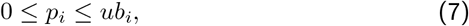

where *m* · *p* determines the inferred mass shift. We can explore the solution space and solve for the next optimal solution by setting a minimal difference *ϵ*_*min*_, or mass error between inferred and observed mass shift.

#### 2.2.3 Linear program that combines both objective functions

In order to find the PTM pattern with the smallest number of PTMs that also minimizes the error between observed and inferred mass shift, we defined an objective function that sums both previously mentioned functions together:

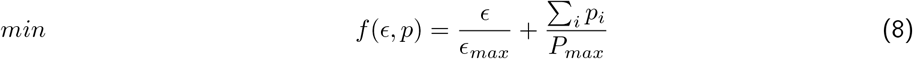

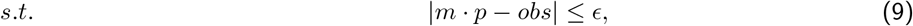

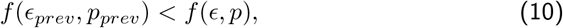

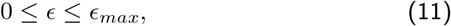

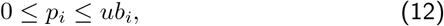

the mass difference *ϵ* and number of PTMs are variables of the LP and are normalized before adding them up together in function *f* (*ϵ, p*). Again, we use the previous value, here denoted as *f* (*ϵ*_*prev*_, *p*_*prev*_), to constrain the solution space and find the next optimal solution.

### 2.3 I^2^MS data of endogenous p53 extracted from MCF7 cells

Experimental I^2^MS data were generated for endogenous p53, which was extracted from MCF7 cells by immunoprecipitation. Two conditions were used to obtain endogenous p53, which is expressed at low levels under basal conditions. In the first condition, p53 levels were enhanced by adding 10 *μ*M Nutlin-3A, which inhibits the E3 ubiquitin ligase MDM2 and thereby prevents p53 degradation. Nutlin-3a does not affect p53 PTMs. The second condition used 10 *J/m*^2^ ultraviolet (UV) radiation to induce DNA damage in cells, which in turn activates kinases known to phosphorylate p53 (e.g. ATR, CHK1) and create further modifications (Liu et al., 2019). Samples were taken 7.5 hours after treatment, at which time p53 levels were elevated (Supplementary Fig. S3). For each condition, two replicates are available.

### 2.4 Generation of mass spectra from simulated PTM pattern distributions

We generated mass spectra from simulated PTM pattern distributions in order to evaluate the algorithm’s performance. The PTM pattern distributions are generated manually and given in the Supplementary Table S1-S3. They range from simple PTM pattern distributions containing only phosphorylation to complex PTM patterns whose isotopic distributions are overlapping. The isotopic distributions are calculated for each PTM pattern using the Python package *pyopenms*. To make the simulated data more realistic, we added noise and error estimated from the experimental I^2^MS data described in the previous section.

We added different types of noise and error to the theoretical isotopic distributions: (i) basal noise, (ii) vertical error, and (iii) horizontal error. An isotopic distribution without error or noise follows approximately a Gaussian distribution and the peaks have a spacing of 1 Da, see Supplementary Fig. S2 for an example of unmodified p53. To estimate the basal noise, a region within the mass spectrum without any signal was selected, from 47 kDa to 49 kDa (Supplementary Fig. S4B and S5A). The horizontal error is defined as any deviation from the standard spacing of 1 Da between two peaks (Supplementary Fig. S5B). For the vertical error, a Gaussian model was fit to the region where the signal is located that presumably contains the modforms of p53 (Supplementary Fig. S4A). The ratio between the fit and the respective peaks is plotted as vertical error in Supplementary Fig. S5C. We fit beta distributions to the noise and error distributions depicted in Supplementary Fig. S5 and randomly selected from the fitted distributions to add basal noise and horizontal and vertical error to the theoretical isotopic distributions.

## 3 Results

The algorithm developed in this study is used to infer PTM patterns for a protein of interest from true mass spectra generated by I^2^MS. The algorithm is tested on experimental data from the tumor suppressor protein p53 which has over 100 possible modification sites and is therefore a good example to test the limits of our tool. For the experimental data, we are not able to evaluate the algorithm’s predictions as we do not know the correct PTM patterns. Therefore, we created simulated data sets with manually generated PTM patterns to test the performance of MSModDetector and to indicate the challenges for detecting mass shifts and predicting correct PTM patterns from I^2^MS data.

### 3.1 Challenges of mass shift detection and PTM pattern inference

We are using a simulated data set to understand what properties of the mass spectrum make it difficult to detect mass shifts and infer PTM patterns. In the following sections, we will discuss these properties and analyze their impact on the predictions by using simulated data of overlapping isotopic distributions (Fig. 3) and of phosphorylation patterns on p53 (Fig. 4). The simulated PTM patterns are listed in Supplementary Tables S1 and S2. Noise and error are added to the theoretical spectra and drawn randomly from the distributions estimated from the experimental I^2^MS data (Supplementary Fig. S5). Since noise and error are randomly drawn, we generate each mass spectra a 100 times, so every performance evaluation on simulated data with noise and error is the average of 100 simulations.

**Figure 3:**
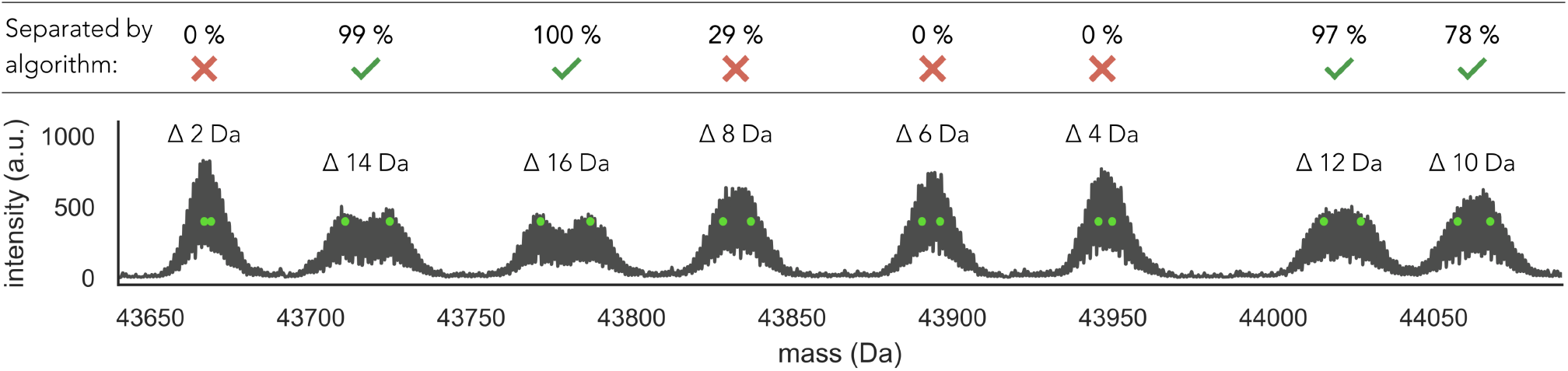
Simulated isotopic distributions with varying amounts of overlap. Basal noise as well as vertical and horizontal error are added to the mass spectrum, which is generated 100 times. Percentages indicate how many times out of the 100 simulations the overlapping distributions could be separated by MSModDetector. Isotopic distributions that overlap by less than 10 Da cannot be separated by the algorithm.

**Figure 4:**
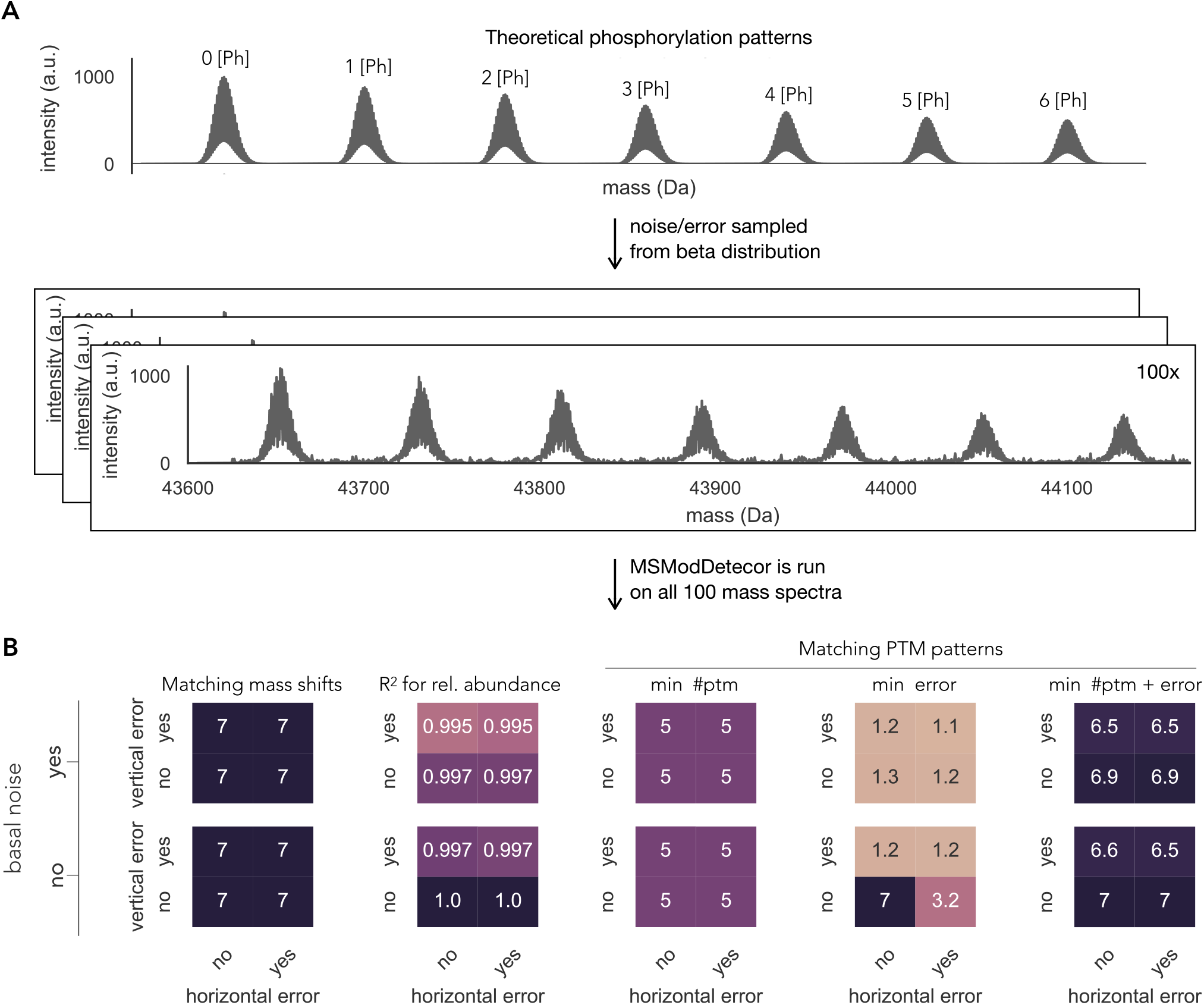
Impact of noise and error on the algorithm’s prediction. (A) Theoretical mass spectrum of manually generated p53 phosphorylation patterns. Basal noise, horizontal and vertical error are sampled randomly 100 times from the respective beta distributions (see Supplementary Fig. S5). (B) Performance evaluation of the 7 mass shifts shown in (A). The mass tolerance is set to 20 ppm and each value depicted here is the average of 100 simulations. On the left, the number of detected mass shifts is displayed for all different combinations of noise and error, and how well their predicted abundances match the observed abundances is shown. On the right, the average values for all correct PTM pattern predictions are shown for three different objective functions.

#### 3.1.1 Overlapping isotopic distributions

Isotopic distributions of protein molecules with different masses can overlap in the mass spectrum and make the task of resolving adjacent distributions challenging. In our example we only look at a selected protein of interest, hence the different masses are caused by different PTM patterns. A phosphorylation for instance adds ~80 Da to the protein’s mass while 2 acetylations add ~ 84 Da to the protein’s mass. While this is no problem for the linear program to distinguish between these two PTM patterns, it is challenging to differentiate between both isotopic distributions. The closer the distributions are, the more likely the algorithm will detect only one distribution with a skewed mass shift value that lies in between both distributions. To test how much overlap can be tolerated by the algorithm in order to be able to separate adjacent distributions, we generated a data set with decreasing amounts of overlaps. Fig. 3 shows isotopic distributions that differ in their average mass value by 2 Da to 16 Da. We added basal noise and vertical and horizontal error to the distributions and ran 100 simulations. Two isotopic distributions are assumed to be successfully separated by MSModDetector if the mean values of both distributions could be detected within a mass tolerance of 20 ppm in more than 75% of the simulations. Adjacent isotopic distributions whose mean values differ by at least 10 Da can be separated by MSModDetector. A difference in 8 Da could only be separated in 29% of the times and everything less than 8 Da could not be separated.

#### 3.1.2 Basal noise, horizontal and vertical error

Next, we analyzed the impact of basal noise as well as horizontal and vertical error, individually and in combination, on mass shift detection using manually generated phosphorylation patterns on p53 (Fig. 4A). The underlying phosphorylation pattern data are shown in Supplementary Table S2. Every performance evaluation in Fig. 4B is based on 100 simulations of the mass spectrum with the respective noise/error and the depicted values indicate the average value of 100 predictions.

Experimentally obtained mass spectra contain basal noise which are fluctuations of the background level in the absence of an analyte. Therefore we need a threshold to distinguish between signal and noise. By choosing a high threshold, we might loose PTM patterns that are of low abundance; but with a low threshold, the algorithm might detect an increased number of incorrect mass shifts. In the example of multiple phosphorylations in Fig. 4B, adding basal noise to the clean theoretical mass spectrum does not influence mass shift detection. MSModDetector also predicts the relative abundance of each detected mass shift. We indicate the accuracy of the prediction by calculating the R-squared value between the observed and predicted relative abundance. The relative abundance prediction is only slightly altered when the data contains basal noise (Fig. 4B).

In addition to basal noise, experimentally obtained I^2^MS spectra show error/variation in the horizontal and vertical axes. For the phosphorylation pattern example given in Fig. 4A, the algorithm is robust against these errors in terms of mass shift detection and only slightly impacted by vertical error for estimating abundances (Fig. 4B).

#### 3.1.3 Impact of the objective function on PTM pattern prediction

PTM pattern prediction is mostly impacted by the choice of an objective function for the linear program. To test the performance of different objective functions, we used the detected mass shifts described in the previous section and predicted PTM patterns for them using three different objective functions: minimizing the number of total PTMs (min #ptm), minimizing the error between observed and inferred mass shifts (min error), and minimizing both the number of PTMs and the error (min #ptm + error). For more details on the different objective functions and linear programs see section 2.2 in Methods.

The mass tolerance for this evaluation, i.e. maximum allowed error between observed and inferred mass shift, is set to 20 ppm. We observe that minimizing the error between inferred and observed mass shift is only a good choice if there is no basal noise or horizontal/vertical error (Fig. 4B). If we choose the objective function that minimizes the number of PTMs, we are not able to predict correct PTM patterns for the two highest mass shifts (∼ 400 Da and ∼480 Da). The bigger the mass shift is, the more combinations are possible for the respective mass and the more difficult it becomes to infer the correct PTM pattern. Furthermore, increasing the mass tolerance expands the number of feasible solutions, and also poses a greater challenge for the algorithm to infer the correct PTM pattern (Supplementary Fig. S6). The combination of both objectives, minimizing error and the number of PTMs, gives the best results (Fig. 4B). However, we want to emphasize that minimizing the number of PTMs is computationally faster.

### 3.2 Evaluation on a complex PTM pattern landscape

As a next step, we generated a complex PTM pattern data set (Fig. 5A) with 4 different PTM types (phosphorylation, acetylation, cysteinylation, oxidation) to further test the algorithm’s performance on a simulated data set that visually resembles the experimental data of endogenous p53, which will be analyzed in the next section. The mass tolerance is set to 36 ppm for this evaluation to see how well MSModDetector performs when predicting PTM patterns given a higher mass error allowance. The PTM patterns of the complex data set are listed in the Supplementary Table S3. Basal noise and error are added to the theoretical mass spectrum, which is then generated 100 times. We tested the objective function that minimizes the number of PTMs as well as the objective function that minimizes both the number of PTMs and the error between inferred and observed mass shifts. Fig. 5B shows for each mass shift how often they could be detected out of 100 simulations and how many of the PTM pattern predictions are correct for both objective functions. A PTM pattern prediction is correct if it matches the PTM pattern in the simulated data set. Furthermore, a mass shift is considered to be successfully detected if it is detected in more than 75% of simulations. The same threshold applies to PTM pattern predictions.

**Figure 5:**
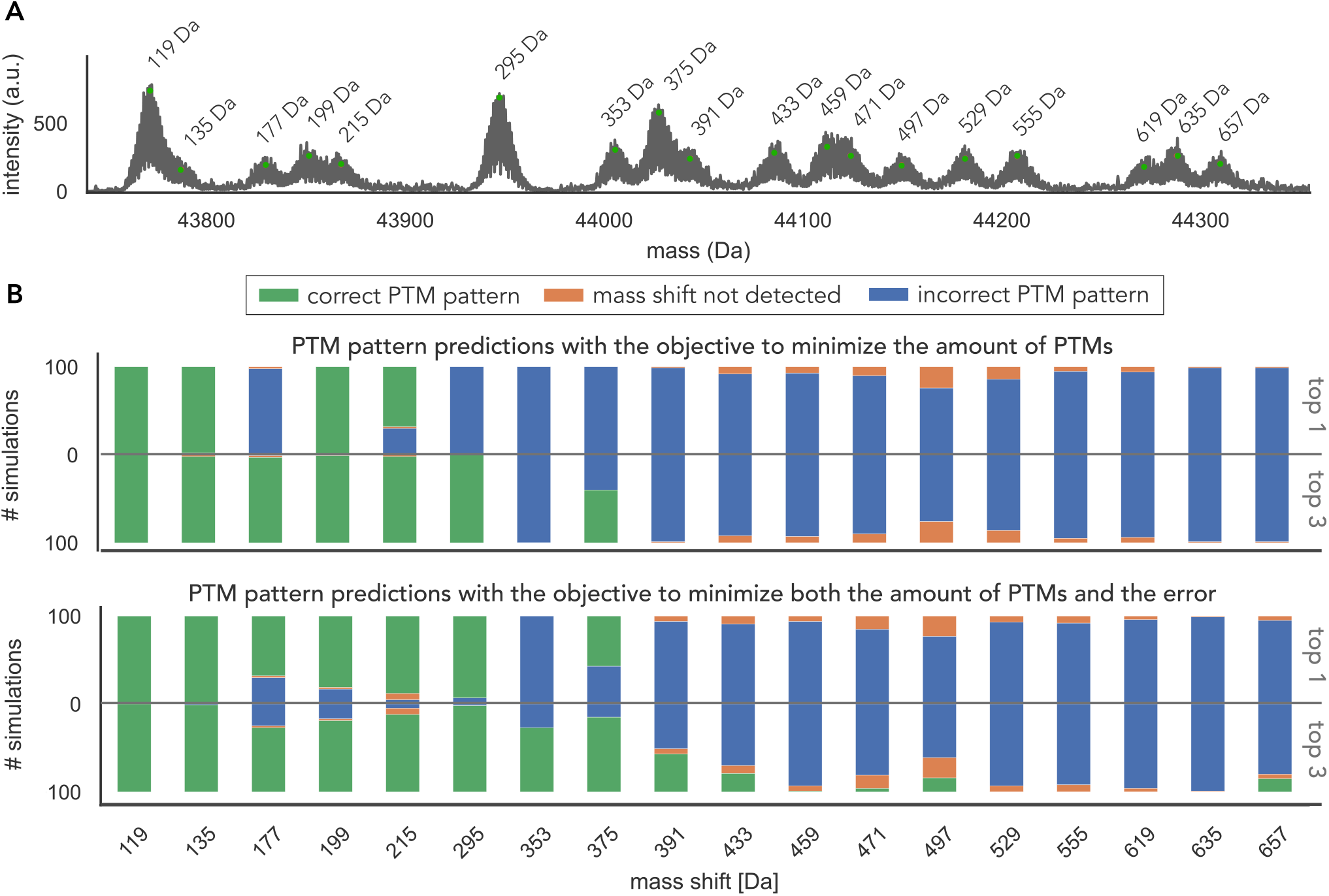
Evaluation of PTM pattern predictions for a complex PTM pattern landscape. (A) Basal noise and vertical and horizontal error are added to the complex PTM pattern spectrum. The respective PTM patterns are listed in the Supplementary Table S3. (B) The mass spectrum is generated 100 times, once to evaluate MSModDetector using the objective function to minimize the number of PTMs, and once using the objective to minimize both the number of PTMs and the error between inferred and observed mass shift. For each objective function, we show how often the correct PTM pattern is predicted considering only the first optimal solution or the 3 optimal solutions.

The performance for this complex PTM pattern landscape is notably reduced compared to the simulated data with phosphorylation patterns only. All 18 mass shifts could be detected and 3 PTM patterns are predicted correctly using the objective that minimizes the number of PTMs. For the objective that minimizes both mass error and number of PTMs, 5 PTM patterns are predicted correctly. By further iterating through the solution space and reporting the 3 optimal solutions, instead of only the first optimal solution, MSModDetector predicts 6 correct PTM patterns with both objective functions (Fig. 5B). Considering the 10 optimal solutions increases the number of correct predictions to 7 (Supplementary Fig. S7 and S8), however, in general there was minimal benefit to executing more than 3 iterations, and increased iterations run the risk of detecting more incorrect solutions.

The PTM patterns for the mass shifts 177 Da and 353 Da contain acetylation, which has approximately the same mass as tri-methylation (∼ 42 Da) and makes it therefore difficult to resolve. Furthermore, incorrect PTM pattern predictions increase with higher mass shifts as more combinations become possible making it more challenging for the algorithm to find the correct PTM pattern. We observe that from the mass shift of 391 Da MSModDetector is not able to predict correct PTM patterns.

Taken together, these results show that by iterating through the solution space and by looking at the combined predictions of both objective functions, we obtain the correct PTM patterns for the first 8 lowest mass shifts up to 391 Da.

### 3.3 Detecting mass shifts in experimental I^2^MS data of endogenous p53

MSModDetector is tested on experimental I^2^MS data of endogenous p53 extracted from MCF7 cells treated with Nutlin-3A or UV as described in section 2.3. Fig. 6 shows the p53 signal that is located in the mass region between 43750 Da and 44520 Da of the I^2^MS spectrum. The p53 signal exhibits multiple isotopic distributions, indicating a complex PTM pattern landscape. The algorithm detects 30 mass shifts across all conditions and replicates, represented by black dots in Fig. 6. Each mass shift is determined as the mean value of the respective isotopic distribution to which it was fit to. The maximum mass shift is around 800 Da.

**Figure 6:**
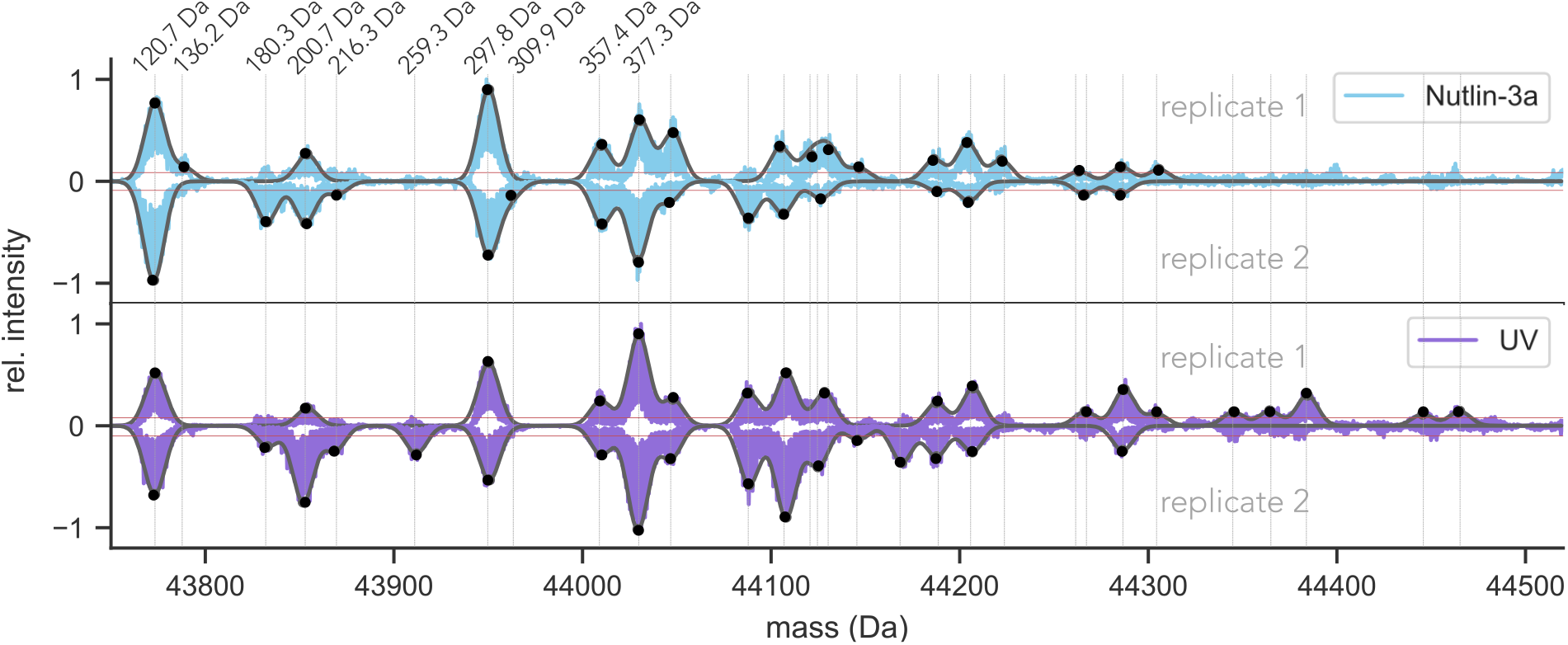
Analysis of endogenous p53 PTM patterns in I MS data. (A) True mass spectra of endogenous p53 extracted from MCF7 cells. Two conditions (cells supplemented with Nutlin-3a or perturbed with UV irradiation) with two replicates each are shown. Red horizontal lines indicate noise levels. The vertical gray lines mark the detected mass shifts. Potential PTM patterns for mass shifts up to 377.3 Da are listed in Table 1. The 3 optimal PTM patterns for all detected mass shift are listed in Supplementary Table S4.

The combinations of possible PTM patterns increase for bigger mass shifts. Using a mass tolerance of 36 ppm, we observe that for a mass shift of approximately 400 Da, around 2500 potential combinations of PTM patterns are possible (see Supplementary Fig. S9) when considering the 9 different PTM types mentioned in Table 1. As the potential PTM pattern increases for higher mass shifts, the problem of PTM pattern inference becomes more challenging. Since the previous evaluation showed that PTM pattern predictions for mass shifts above 400 Da are not correct, we only list potential PTM patterns for detected mass shifts under 377.3 Da in Table 1. Table 1 lists the optimal PTM patterns for two different objective functions. Potential PTM patterns for all other detected mass shifts are listed in the Supplementary Table S4.

## 4 Conclusions

We developed a Python module to process I^2^MS data and study PTM patterns of a protein of interest. The module consists of an algorithm that detects mass shifts and a linear program to infer potential PTM patterns. The algorithm was tested on experimental data of endogenous p53 and on simulated data to put the PTM pattern predictions in context and get an estimate how many predictions are correct on average.

Using I^2^MS based information reveals the maximum mass shift of a protein. Surprisingly, the analysis of I^2^MS of p53 shows that, at this level of detection, the protein has only a maximal mass shift of approximately 800 Da. Even though p53 has more than 100 modification sites, only a few of them seem to be occupied at the same time on a given individual p53 protein.

Noise and error in the data only slightly impacts PTM pattern predictions, whereas the choice of the objective function has a big influence. Furthermore, overlapping distributions with average masses differing by less than 10 Da cannot be separated by the algorithm and skew the mass shift detection, leading to incorrect PTM pattern predictions.

We tested the algorithm’s performance on a complex PTM pattern data set and observed that the PTM patterns of mass shifts higher than approximately 400 Da cannot be resolved. Iterating through the solution space can increase the number of correct PTM pattern predictions. The user can specify how many iterations or rounds of optimizations for each mass shift should be done by MSModDetector. The tool will list the best k solutions, however, the user has no information on which of the best k optimal solutions is the correct PTM pattern or if the correct PTM pattern is included in the k optimal solutions. For our simulated example we did not observe significant improvements after 3 iterations (see Supplementary Fig. S7 and S8). Therefore, we listed the top 3 optimal PTM pattern predictions for the experimental I^2^MS data of endogenous p53 in Supplementary Table S4 for two objective functions.

Whilst MSModDetector is useful for comparing PTM pattern landscapes across samples, MS^1^ based analysis is not able to localize the PTMs, for that task MS^2^ data is required, a process we are currently developing. Once MS^2^ data is easily accessible for the I^2^MS approach, MSModDetector should be extended to incorporate MS^2^ information for PTM localization.

Taken together, this Python module is a useful tool to get first insights into a protein’s PTM pattern landscape and helps to compare it across samples to identify significant differences that might give rise to distinct biological outcomes. The tool can be used as an initial analysis method to highlight important differences between PTM pattern landscapes of a protein of interest in different samples. Interesting mass shifts and their potential PTM pattern predictions can then be further validated or studied by other experimental methods. These insights will have important implications for the role of PTM patterns in disease processes and will help to determine therapeutic targets.

## Acknowledgements

We thank Michael A. R. Hollas and Bryan P. Early for helpful comments on the MSModDetector code and on the draft.

## Data availability

The data underlying this article are available in MassIVE and and can be accessed with MSV000091481. It was assigned doi:10.25345/C55Q4RW7Z.

## References

R Aebersold et al. How many human proteoforms are there? Nature Chemical Biology, 14(3):206–214, 2018.

W Cai, H Guner, ZR Gregorich, AJ Chen, S Ayaz-Guner, Y Peng, SG Valeja, X Liu, and Y Ge. Mash suite pro: A comprehensive software tool for top-down proteomics. Molecular & Cellular Proteomics, 15(2):703–14, 2016.

Philip D. Compton, Neil L. Kelleher, and Jeremy Gunawardena. Estimating the distribution of protein post-translational modification states by mass spectrometry. Journal of Proteome Research, 17(8):2727–2734, 2018.

Veronika Csizmok and Julie D Forman-Kay. Complex regulatory mechanisms mediated by the interplay of multiple post-translational modifications. Current Opinion in Structural Biology, 48:58–67, 2018.

Antonina Hafner, Martha L Bulyk, Ashwini Jambhekar, and Galit Lahav. The multiple mechanisms that regulate p53 activity and cell fate. Nature Reviews Molecular Cell Biology, 20(4):199–210, 2019.

DM Horn, RA Zubarev, and FW McLafferty. Automated reduction and interpretation of high resolution electrospray mass spectra of large molecules. The Journal of the American Society for Mass Spectrometry, 11(4):320–32, 2000.

Q Huang, J Chang, MK Cheung, W Nong, L Li, MT Lee, and Kwan HS. Human proteins with target sites of multiple post-translational modification types are more prone to be involved in disease. Journal of Proteome Research, 13 (6):2735–2748, 2014.

O Jensen. Interpreting the protein language using proteomics. Nature Reviews Molecular Cell Biology, 7:391–403, 2006.

Kyowon Jeong, Jihyung Kim, Manasi Gaikwad, Siti Nurul Hidayah, Laura Heikaus, Hartmut Schlüter, and Oliver Kohlbacher. Flashdeconv: Ultrafast, high-quality feature deconvolution for top-down proteomicss. Cell Systems, 10 (2):213–218, 2020.

JO Kafader, KR Durbin, RD Melani, BJ Des Soye, LF Schachner, MW Senko, PD Compton, and NL Kelleher. Individual ion mass spectrometry enhances the sensitivity and sequence coverage of top-down mass spectrometry. Journal of Proteome Research, 19(3):1346–1350, 2020.

AT Kong, FV Leprevost, DM Avtonomov, D Mellacheruvu, and AI Nesvizhskii. MSFragger: ultrafast and comprehensive peptide identification in shotgun proteomics. Nature Methods, 14(5):513–520, 2017.

Qiang Kou, Likun Xun, and Xiaowen Liu. TopPIC: a software tool for top-down mass spectrometry-based proteoform identification and characterization. Bioinformatics, 32(22):3495–3497, 2016.

Mario Leutert, Samuel W Entwisle, and Judit Villén. Decoding post-translational modification crosstalk with proteomics. Molecular & Cellular Proteomics, 20(100129):1535–9476, 2021.

X Liu, Y Sirotkin, Y Shen, G Anderson, YS Tsai, YS Ting, DR Goodlett, RD Smith, V Bafna, and PA Pevzner. Protein identification using top-down spectra. Molecular & Cellular Proteomics, 11(6):M111.008524, 2012.

Y Liu, O Tavana, and W Gu. p53 modifications: exquisite decorations of the powerful guardian. Journal of Molecular Cell Biology, 11(7):564–577, 2019.

RD Melani, BJ Des Soye, JO Kafader, E Forte, M Hollas, V Blagojevic, F Negrao, JP McGee, B Drown, C Lloyd-Jones, HS Seckler, JM Camarillo, PD Compton, RD LeDuc, B Early, RT Fellers, BK Cho, BB Mattamana, YA Goo, PM Thomas, MK Ash, PP Bhimalli, L Al-Harthi, BE Sha, JR Schneider, and NL Kelleher. Next-generation serology by mass spectrometry: Readout of the sars-cov-2 antibody repertoire. Journal of Molecular Cell Biology, 21(1): 274–288, 2022.

Sudhakaran Prabakaran, Guy Lippens, Hanno Steen, and Jeremy Gunawardena. Post-translational modification: Nature’s escape from genetic imprisonment and the basis for dynamic information encoding. Wiley Interdisciplinary Reviews: Systems Biology and Medicine, 4(6):565–583, 2012.

LV Schaffer, RJ Millikin, RM Miller, LC Anderson, RT Fellers, Y Ge, NL Kelleher, RD LeDuc, X Liu, SH Payne, L Sun, PM Thomas, T Tucholski, Z Wang, S Wu, Z Wu, D Yu, MR Shortreed, and LM Smith. Identification and quantification of proteoforms by mass spectrometry. Proteomics, 19(10):e1800361, 2019.

S Tyanova, T Temu, and Cox J. The maxquant computational platform for mass spectrometry-based shotgun proteomics. Nature Protocols, 11:2301–2319, 2016.

