## Supplement for "MSModDetector: A Tool for Detecting Mass Shifts and Post-Translational Modifications in Individual Ion Mass Spectrometry Data"

### Supplemental Figures

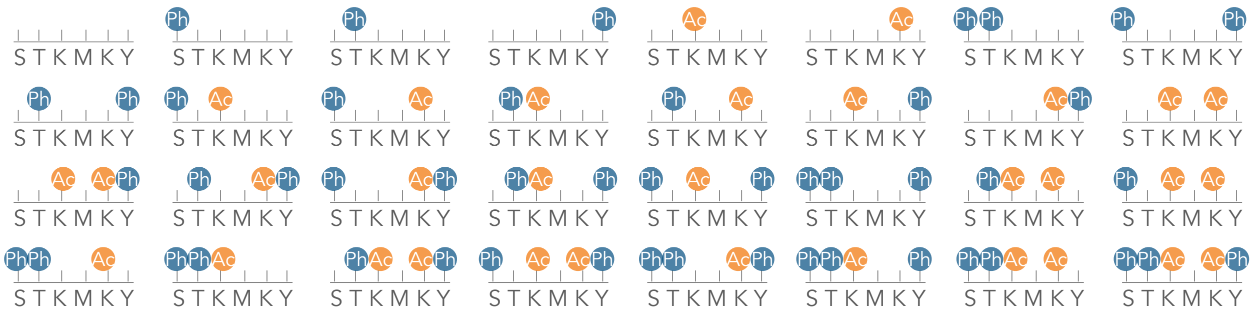

**Figure S1: Combinatorial patterns of PTMs (modforms).** A hypothetical protein harboring 5 modification sites. The modification sites serine (S), threonine (T), and tyrosine (Y) can be phosphorylated (Ph) and the modification site lysine (K) can be acetylated (Ac). This sums up to 32 modforms in total.

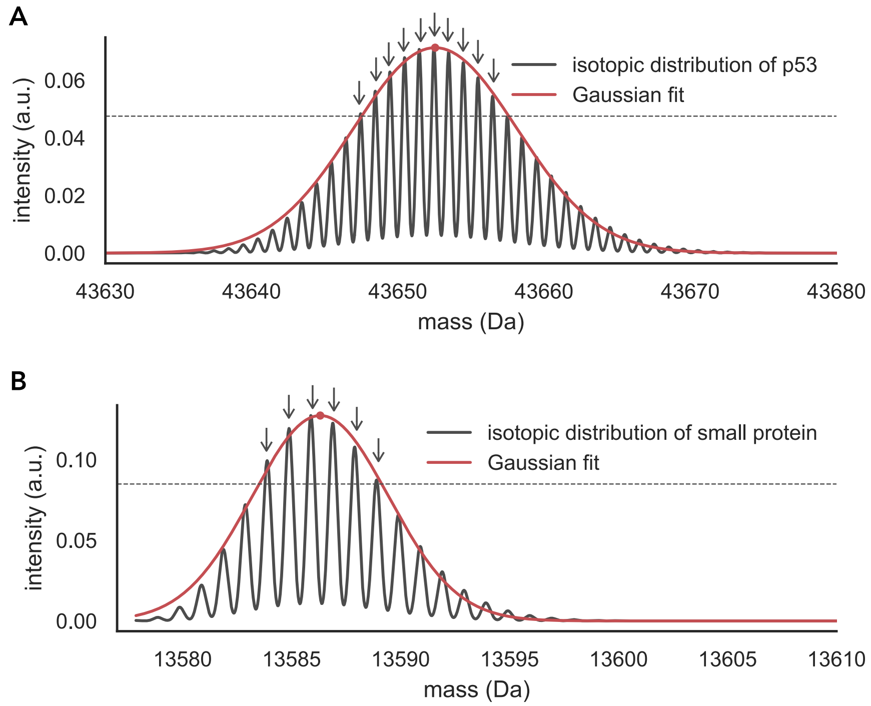


**Figure S2: Gaussian functions fitted to theoretical isotopic distributions of proteins.** Isotopic distributions are simulated using the Python package *pyopenms*. The dotted horizontal line indicates the upper third part of the isotopic distribution and the arrows indicate which peaks are above the dotted line. (A) The Gaussian distribution fitted to unmodified p53 has a mean value of 43652 Da (depicted by the red dot) and the standard deviation is 5.596. 10 peaks are above the dotted line. (B) We chose an arbitrarily small protein (UniProt id: Q9BZX4-2) to showcase its theoretical isotopic distribution. The average mass of this protein is 13587 Da. 6 peaks are above the dotted line.

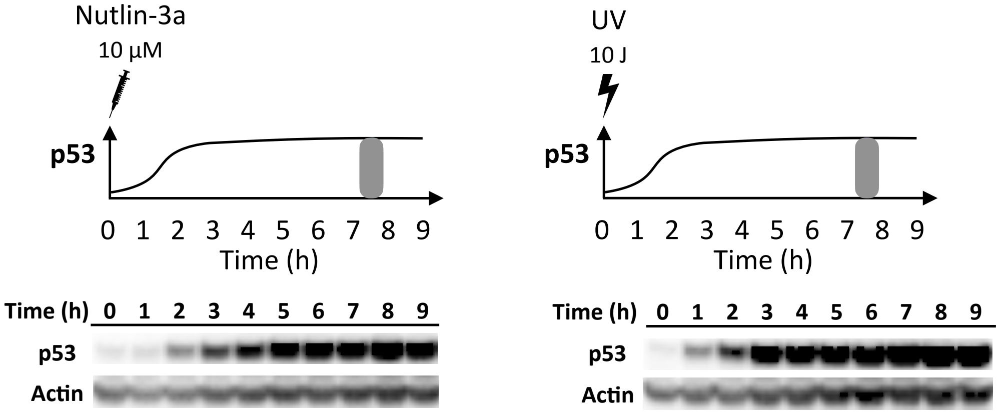


**Figure S3: Western blot for endogenous p53.** MCF7 cells were subjected to either 10 $\mu M$ Nutlin-3a (left panel) or 10 $J/m^2$ UV (right panel) leading to accumulation of p53 levels. Gray bars at 7.5 hours indicate the time point at which samples were taken after treatment.

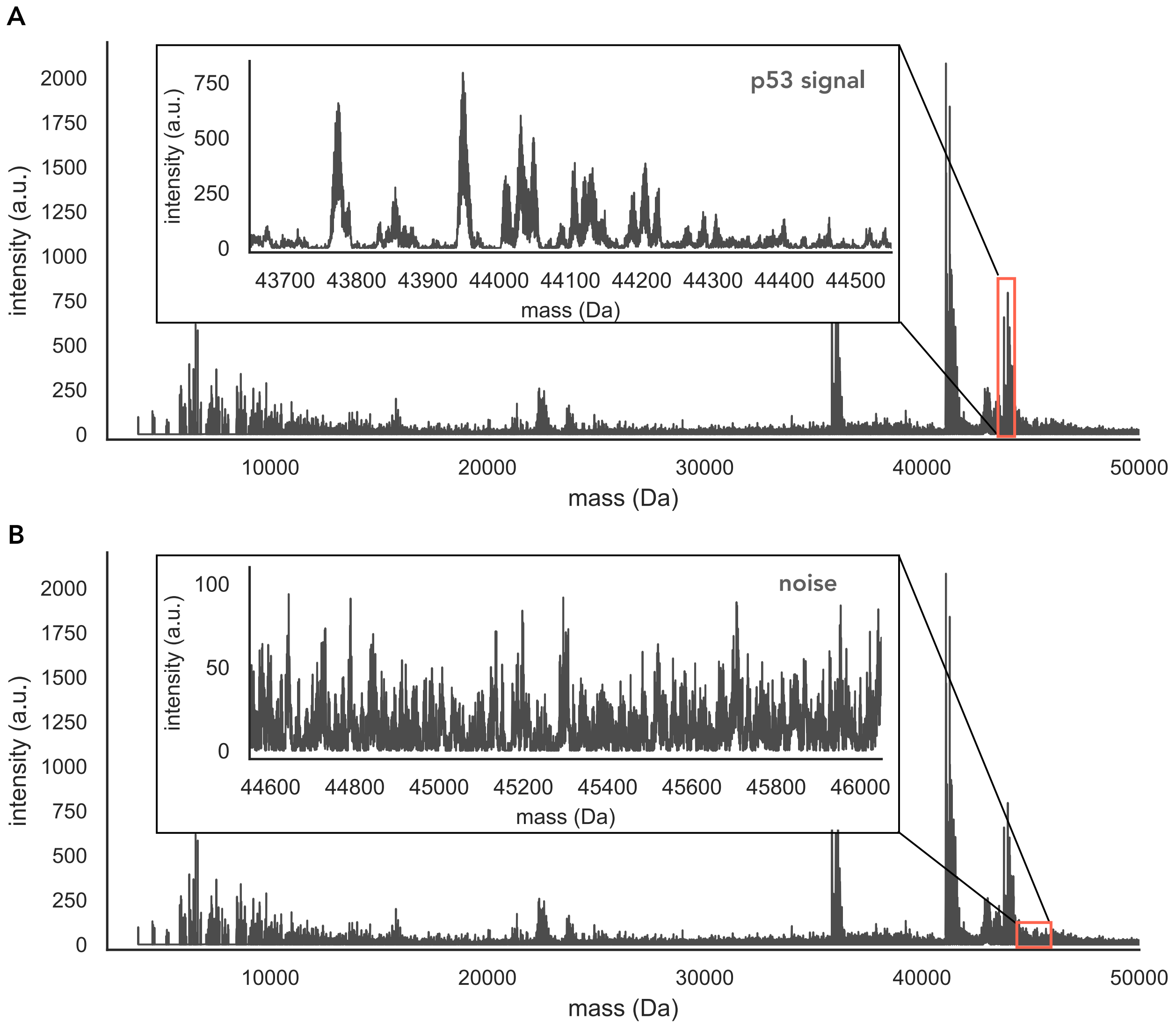


**Figure S4: Mass spectrum example from MCF7 cells supplemented with Nutlin-3a.** p53 was extracted by immunoprecipitation. Insets show (A) the expected mass range for the p53 signal with a monoisotopic mass of 43,653 Da and (B) an example where no signal is observed. This area is used to determine the basal noise in the spectrum.

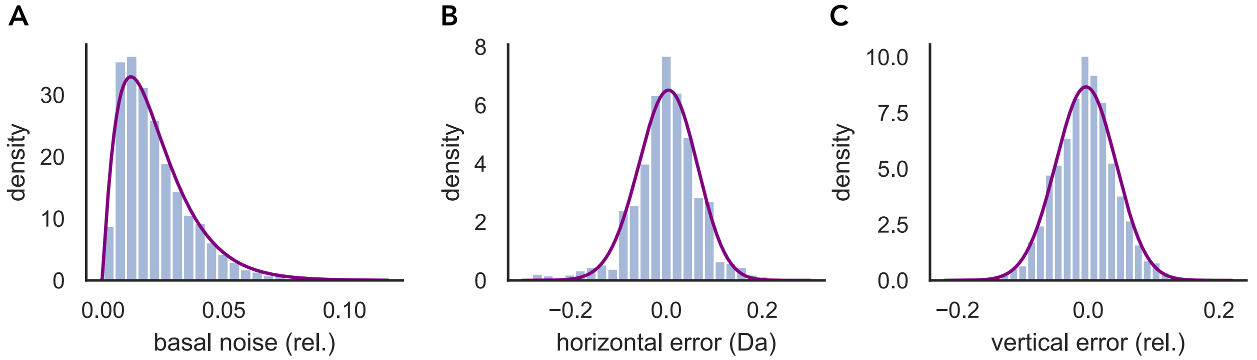


**Figure S5: Noise and error distribution in I$^2$MS data from endogenous p53.** Data was obtained from MCF7 cells under 2 different conditions with two replicates each (Nutlin-3a only and UV radiation). Peaks from a mass range where no signal is observed (44.6kDa - 46kD) is used to obtain the distribution for the basal noise (A). The mass spectrum region between 43.6 kDa and 44.6 kDa is selected to determine horizontal and vertical error (B,C). Mass differences between peaks that differ by 1 Da are considered as horizontal error (B). To calculate the vertical error Gaussian distributions are fitted to the mass spectrum and the relative deviation from the fit to the peaks are used for the distribution of the vertical error. Purple lines represent fitted beta distributions.

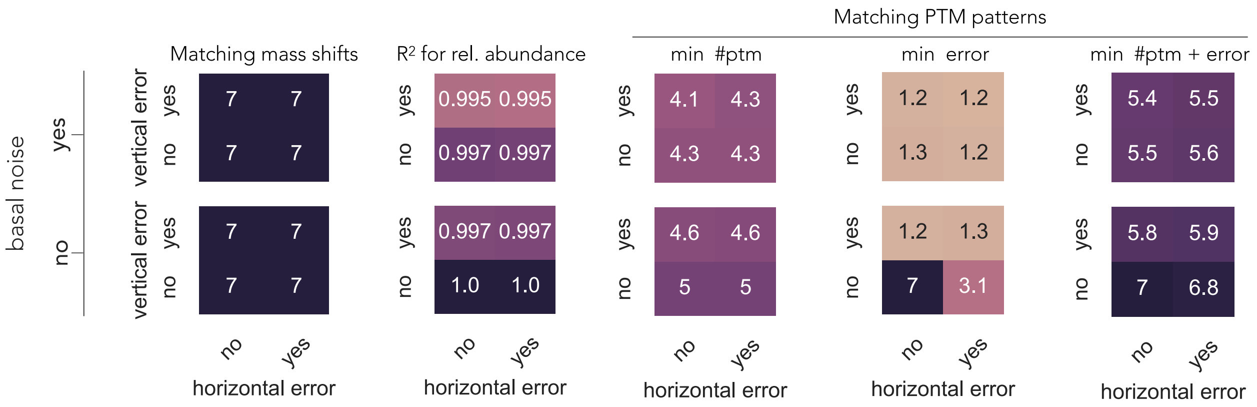


**Figure S6: Impact of noise and error on the algorithm's prediction for phosphorylation patterns on p53.** Theoretical mass spectra of manually generated p53 phosphorylation patterns including basal noise, horizontal and vertical error are generated 100 times. Each value depicted here is the average of 100 simulations. The mass tolerance is set to 36 ppm. The phosphorylation pattern data set contains 7 PTM patterns (see Supplementary Table S1). On the left, the number of detected mass shifts is displayed for all different combinations of noise and error and how well their predicted abundances match the observed abundances. On the right, the average value for all correct PTM pattern predictions is shown for three different objective functions.

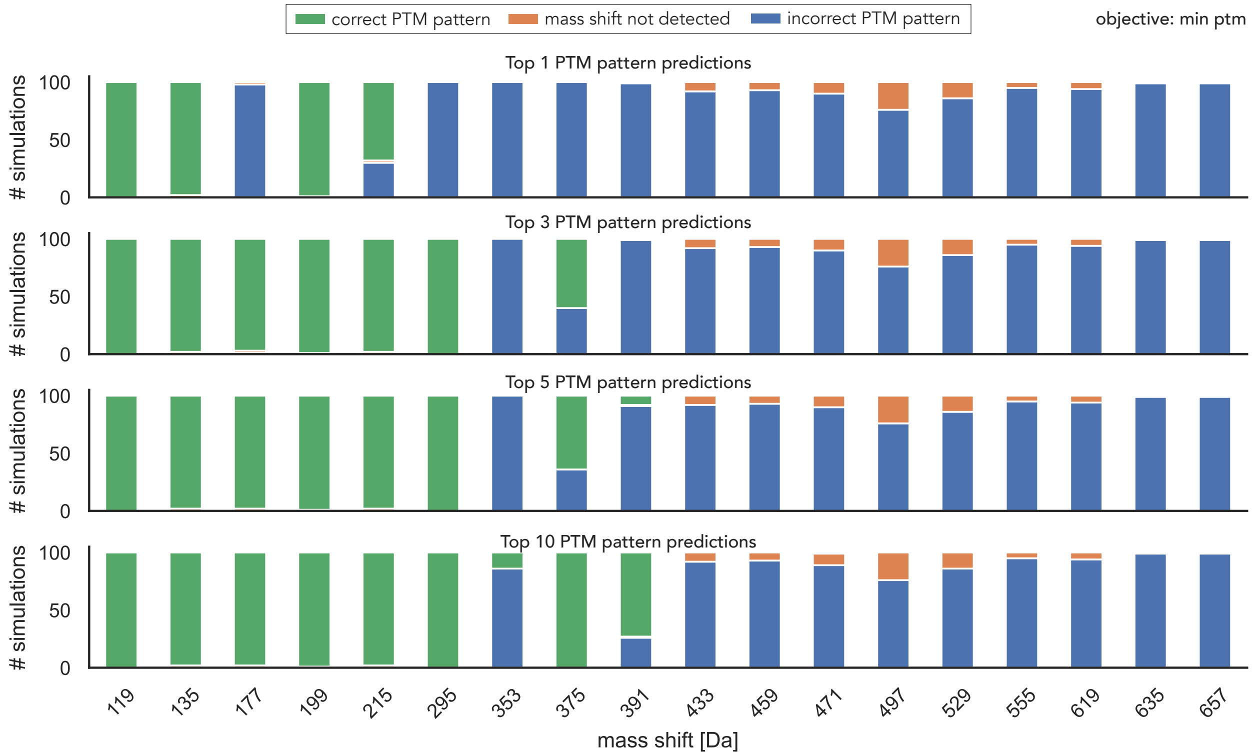


**Figure S7: Evaluation of PTM pattern prediction using the objective to minimize the number of PTMs.** Basal noise and vertical and horizontal error are added to the complex PTM pattern mass spectrum and simulations are run 100 times. The mass tolerance is set to 36 ppm. We observe that iterating through the solution space of the linear program increases the number of correct PTM pattern predictions. If we look at the first optimal solution only (top panel) 3 PTM patterns out of 18 are predicted successfully by MSModDetector. A PTM pattern for a given mass shift is determined to be successfully predicted by MSModDetector if 75%, out of 100 simulations, are predicted correctly. If we consider the 3 optimal solutions for each mass shifts, 6 PTM patterns are successfully predicted. The number of successfully predicted PTM patterns increases to 7 if we consider the 10 optimal solutions.

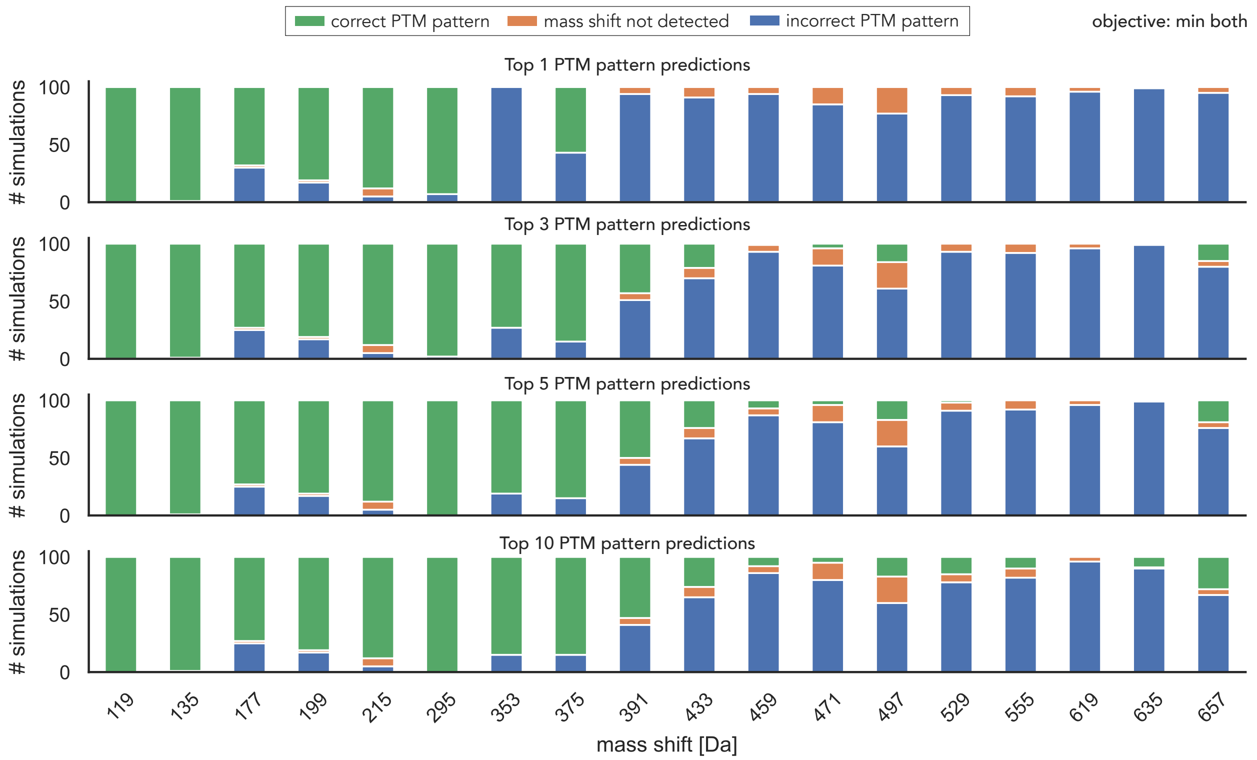


**Figure S8: Evaluation of PTM pattern prediction using the objective to minimize both the number of PTMs and the error between observed and inferred mass shifts.** Basal noise and vertical and horizontal error are added to the complex PTM pattern mass spectrum and simulations are run 100 times. The mass tolerance is set to 36 ppm. As in Supplementary Fig. S6, we observe that iterating through the solution space of the linear program increases the number of correct PTM pattern predictions. If we look at the first optimal solution only (top panel) 5 PTM patterns out of 18 are predicted successfully by MSModDetector. A PTM pattern for a given mass shift is determined to be successfully predicted by MSModDetector if 75%, out of 100 simulations, are predicted correctly. If we consider the 3 optimal solutions for each mass shifts, 6 PTM patterns are successfully predicted. The number of successfully predicted PTM patterns increases to 7 if we consider the top 5 or 10 optimal solutions.

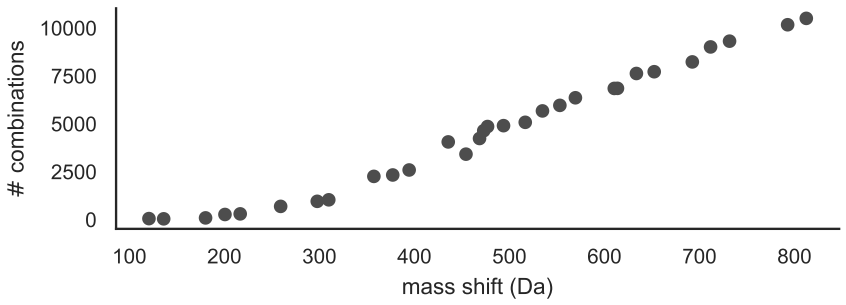


**Figure S9: Potential PTM pattern combinations increase for higher mass shifts.** Mass shifts are obtained from $I^2MS$ measurements of endogenous p53 under Nutlin-3a and UV conditions (see Figure 6 in the main text). For every detected mass shift the linear program with the objective to minimize the number of PTMs is solved and the solution space is explored for a maximum mass tolerance of 36 ppm. The number of possible PTM pattern combinations is shown for each mass shift. The following PTM types are considered for the pattern combinations: phosphorylation (Ph), acetylation (Ac), methylation (Me1), di-methylation (Me2), tri-methylation (Me3), phosphate (Ph-OH), oxidation (Ox), cysteinylation (Cys), sodium adduct (Na).

### Supplemental Tables

**Table S1: Manually generated PTM patterns with different overlaps in their isotopic distributions.** PTM patterns are composed from the following PTM types: phosphorylation (Ph), acetylation (Ac), methylation (Me1), cysteinylation (Cys), and oxidation (Ox).

| index | PTM pattern | average mass (Da) | intensity (a.u.) |
|---|---|---|---|
| 1 | 2[Ph]1[Ox]1[Cys] | 295.1015 | 400 |
| 2 | 3[Ph]2[Ox]1[Na] | 293.92030 | 400 |
| 3 | 1[Me1] | 14.0266 | 400 |
| 4 | 1[Ox] | 15.9994 | 400 |
| 5 | 2[Ph]1[Ac]1[Me1]1[Cys] | 335.1654 | 400 |
| 6 | 1[Ph]3[Ac]1[Me1]1[Cys] | 339.2589 | 400 |
| 7 | 2[Ph]1[Ac]1[Me1]1[Na] | 238.0049 | 400 |
| 8 | 2[Ph]2[Ac] | 244.0332 | 400 |
| 9 | 2[Ph]1[Ox] | 175.9592 | 400 |
| 10 | 4[Ac]1[Ox] | 184.14620 | 400 |
| 11 | 3[Ph]3[Ac]1[Cys] | 485.1921 | 400 |
| 12 | 4[Ph]1[Ac]1[Me1]1[Cys] | 495.1252 | 400 |
| 13 | 3[Ph]2[Ac]1[Cys] | 443.1554 | 400 |
| 14 | 4[Ph]1[Ox]1[Cys] | 455.0613 | 400 |
| 15 | 1[Ac]1[Ox] | 58.0361 | 400 |
| 16 | 1[Ac]1[Me1]1[Ox] | 72.0627 | 400 |
| 17 | 1[Cys] | 119.1423 | 400 |
| 18 | 1[Ox]1[Cys] | 135.1417 | 400 |

**Table S2: Manually generated phosphorylation patterns on p53.**

| index | PTM pattern | average mass (Da) | intensity (a.u.) |
|---|---|---|---|
| 1 | 0 | 0 | 994 |
| 2 | 1[Ph] | 79.9799 | 875 |
| 3 | 2[Ph] | 159.9598 | 793 |
| 4 | 3[Ph] | 239.9397 | 667 |
| 5 | 4[Ph] | 319.9196 | 592 |
| 6 | 5[Ph] | 399.8995 | 527 |
| 7 | 6[Ph] | 479.8794 | 499 |

**Table S3: Manually generated complex PTM pattern data set.** PTM patterns are composed from the following PTM types: phosphorylation (Ph), acetylation (Ac), cysteinylation (Cys), and oxidation (Ox).

| index | PTM pattern | mass | intensity |
|---|---|---|---|
| 1 | 1[Cys] | 119.1423 | 732 |
| 2 | 1[Cys]1[Ox] | 135.1417 | 162 |
| 3 | 1[Cys]1[Ac]1[Ox] | 177.1784 | 188 |
| 4 | 1[Cys]1[Ph] | 199.1222 | 265 |
| 5 | 1[Cys]1[Ph]1[Ox] | 215.1216 | 191 |
| 6 | 1[Cys]2[Ph]1[Ox] | 295.1015 | 689 |
| 7 | 1[Cys]2[Ph]1[Ac]2[Ox] | 353.1376 | 308 |
| 8 | 1[Cys]3[Ph]1[Ox] | 375.0814 | 577 |
| 9 | 1[Cys]3[Ph]2[Ox] | 391.0808 | 243 |
| 10 | 1[Cys]3[Ph]1[Ac]2[Ox] | 433.1175 | 281 |
| 11 | 1[Cys]3[Ph]2[Ac]1[Ox] | 459.1548 | 326 |
| 12 | 1[Cys]4[Ph]2[Ox] | 471.0607 | 264 |
| 13 | 1[Cys]4[Ph]1[Ac]1[Ox] | 497.098 | 198 |
| 14 | 1[Cys]4[Ph]1[Ac]3[Ox] | 529.0968 | 246 |
| 15 | 1[Cys]4[Ph]2[Ac]2[Ox] | 555.1341 | 251 |
| 16 | 1[Cys]5[Ph]2[Ac]1[Ox] | 619.1146 | 179 |
| 17 | 1[Cys]5[Ph]2[Ac]2[Ox] | 635.114 | 261 |
| 18 | 1[Cys]6[Ph]1[Ac]1[Ox] | 657.0578 | 208 |

**Table S4: Top 3 PTM pattern predictions for the $I^2MS$ data of endogenous p53 under Nutlin-3a and UV conditions.** PTM pattern predictions are shown for two different objective functions: "min both" is minimizing the number of PTMs and the error between observed and inferred mass shift and "min ptm" is only minimizing the number of PTMs. The following PTM types are considered for the pattern combinations: phosphorylation (Ph), acetylation (Ac), methylation (Me1), di-methylation (Me2), tri-methylation (Me3), phosphate (Ph-OH), oxidation (Ox), cysteinylation (Cys), sodium adduct (Na).

| mass shift | PTM pattern (min both) | PTM pattern (min ptm) |
|---|---|---|
| 120.69 | 1[Ph-OH]1[Na] | 1[Cys] |
| 120.69 | 1[Ph]1[Ac] | 1[Ph-OH]1[Na] |
| 120.69 | 1[Cys] | 1[Ph]1[Me3] |
| 136.2 | 1[Ph]2[Me2] | 1[Cys]1[Ox] |
| 136.2 | 1[Ph]1[Ac]1[Me1] | 1[Ph]2[Me2] |
| 136.2 | 1[Ph-OH]1[Na]1[Ox] | 1[Ph]1[Me3]1[Me1] |
| 180.28 | 1[Ph]2[Me3]1[Ox] | 1[Ph]2[Me3]1[Ox] |
| 180.28 | 1[Ph]1[Me3]1[Ac]1[Ox] | 1[Ph]1[Me3]1[Ac]1[Ox] |
| 180.28 | 1[Ph]2[Ac]1[Ox] | 1[Cys]2[Na]1[Ox] |
| 200.69 | 1[Ph-OH]1[Ph]1[Na] | 1[Cys]1[Ph] |
| 200.69 | 4[Me3]2[Ox] | 1[Ph-OH]1[Ph]1[Na] |
| 200.69 | 3[Me3]1[Ac]2[Ox] | 2[Ph]1[Me3] |
| 216.74 | 1[Cys]1[Ph-OH] | 1[Cys]1[Ph-OH] |
| 216.74 | 1[Cys]2[Ac]1[Me1] | 2[Ph-OH]1[Na] |
| 216.74 | 1[Cys]1[Me3]1[Ac]1[Me1] | 1[Cys]1[Ac]2[Me2] |
| 259.31 | 1[Cys]1[Ph-OH]1[Me3] | 1[Cys]1[Ph-OH]1[Me3] |
| 259.31 | 1[Cys]1[Ph-OH]1[Ac] | 1[Cys]1[Ph-OH]1[Ac] |

| mass shift | PTM pattern (min both) | PTM pattern (min ptm) |
|---|---|---|
| 259.31 | 1[Cys]1[Ph-OH]1[Me2]1[Me1] | 2[Cys]1[Na] |
| 297.83 | 2[Ph-OH]1[Ph]1[Na] | 1[Cys]1[Ph-OH]1[Ph] |
| 297.83 | 3[Ph]1[Ac]1[Ox] | 2[Ph-OH]1[Ph]1[Na] |
| 297.83 | 1[Cys]1[Ph]1[Me3]2[Me2] | 2[Cys]1[Me3]1[Ox] |
| 309.94 | 3[Ph-OH]1[Ox] | 3[Ph-OH]1[Ox] |
| 309.94 | 3[Ph]1[Ac]1[Me2] | 1[Cys]1[Ph-OH]1[Ph]1[Me1] |
| 309.94 | 1[Ph-OH]2[Ph]1[Na]1[Ox]1[Me1] | 2[Cys]1[Me3]1[Me2] |
| 357.41 | 3[Cys] | 3[Cys] |
| 357.41 | 1[Cys]1[Ph-OH]2[Me3]2[Me2] | 1[Cys]2[Ph-OH]1[Me3] |
| 357.41 | 1[Cys]1[Ph-OH]3[Me3]1[Me1] | 2[Cys]1[Ph-OH]1[Na] |
| 377.28 | 1[Cys]2[Ph]2[Me3]1[Me1] | 1[Cys]1[Ph-OH]2[Ph] |
| 377.28 | 1[Cys]2[Ph]1[Ac]2[Me2] | 2[Cys]1[Ph-OH]1[Ac] |
| 377.28 | 1[Cys]1[Ph-OH]2[Ph] | 2[Cys]1[Ph-OH]1[Me3] |
| 394.55 | 2[Cys]2[Me3]2[Me2]1[Ox] | 1[Cys]2[Ph-OH]1[Ph] |
| 394.55 | 2[Cys]3[Me3]1[Ox]1[Me1] | 2[Cys]1[Ph-OH]1[Me3]1[Ox] |
| 394.55 | 2[Cys]1[Me3]1[Ac]2[Me2]1[Ox] | 3[Cys]1[Na]1[Me1] |
| 435.69 | 2[Ph-OH]3[Ph] | 2[Cys]2[Ph-OH] |
| 435.69 | 4[Ph-OH]2[Na] | 2[Ph-OH]3[Ph] |
| 435.69 | 5[Ph]1[Na]1[Me1] | 1[Cys]2[Ph-OH]1[Ph]1[Ac] |
| 454.36 | 2[Cys]2[Ph]2[Me2] | 3[Cys]1[Ph-OH] |
| 454.36 | 2[Cys]2[Ph]1[Me3]1[Me1] | 3[Ph-OH]2[Ph] |
| 454.36 | 2[Cys]2[Ph]1[Ac]1[Me1] | 1[Cys]3[Ph-OH]1[Ac] |
| 468.62 | 2[Cys]2[Ph]1[Me3]1[Me2] | 3[Cys]1[Ph-OH]1[Me1] |
| 468.62 | 2[Cys]4[Me3]3[Ox]1[Me1] | 2[Cys]2[Ph]1[Me3]1[Me2] |
| 468.62 | 2[Cys]3[Me3]2[Me2]3[Ox] | 3[Cys]1[Ph]2[Ox] |
| 473.05 | 1[Cys]1[Ph-OH]3[Ph]1[Ox] | 4[Ph-OH]1[Ph] |
| 473.05 | 1[Cys]3[Ph-OH]2[Na]1[Ox] | 1[Cys]1[Ph-OH]3[Ph]1[Ox] |
| 473.05 | 1[Cys]2[Ph-OH]1[Ph]1[Ac]1[Na]1[Me1] | 2[Cys]1[Ph-OH]1[Ph]1[Me3]1[Ox] |
| 477.06 | 1[Cys]3[Ph-OH]1[Ac]1[Na] | 4[Cys] |
| 477.06 | 1[Cys]3[Ph-OH]1[Me3]1[Na] | 3[Cys]1[Ph-OH]1[Na] |
| 477.06 | 1[Cys]1[Ph-OH]2[Ph]2[Ac]1[Ox] | 2[Cys]2[Ph-OH]1[Me3] |
| 493.87 | 4[Ph-OH]1[Ph]1[Na] | 1[Cys]3[Ph-OH]1[Ph] |
| 493.87 | 6[Ph]1[Me1] | 4[Cys]1[Ox] |
| 493.87 | 2[Ph-OH]3[Ph]1[Ac]1[Ox] | 3[Cys]1[Ph]1[Me3]1[Me1] |
| 516.74 | 1[Cys]2[Ph-OH]2[Ph]1[Ac] | 3[Cys]2[Ph] |
| 516.74 | 2[Cys]1[Ph]3[Me3]1[Me2]2[Na] | 1[Cys]2[Ph-OH]2[Ph]1[Ac] |
| 516.74 | 1[Cys]1[Ph-OH]3[Ph]2[Na]1[Ox] | 5[Ph-OH]1[Me2] |
| 534.85 | 4[Cys]1[Me3]1[Ox] | 3[Cys]1[Ph-OH]1[Ph] |
| 534.85 | 4[Cys]1[Ac]1[Ox] | 4[Cys]1[Me3]1[Ox] |
| 534.85 | 1[Cys]3[Ph-OH]1[Ph]1[Ac] | 4[Cys]1[Ac]1[Ox] |
| 553.16 | 1[Cys]4[Ph-OH]1[Ac] | 3[Cys]2[Ph-OH] |
| 553.16 | 1[Cys]4[Ph-OH]1[Me3] | 1[Cys]4[Ph-OH]1[Ac] |
| 553.16 | 1[Cys]4[Ph-OH]1[Me2]1[Me1] | 4[Ph-OH]2[Ph] |
| 569.52 | 3[Cys]2[Ph-OH]1[Ox] | 3[Cys]2[Ph-OH]1[Ox] |
| 569.52 | 3[Cys]1[Ph-OH]2[Ac]1[Ox]1[Me1] | 5[Ph-OH]1[Ph] |
| 569.52 | 3[Cys]1[Ph-OH]1[Me3]1[Me2]2[Na] | 4[Cys]1[Ph]1[Me1] |
| 610.44 | 2[Cys]1[Ph-OH]2[Ph]1[Me3]2[Me2]1[Ox] | 5[Cys]1[Me1] |

| mass shift | PTM pattern (min both) | PTM pattern (min ptm) |
|---|---|---|
| 610.44 | 2[Cys]2[Ph-OH]3[Ac]1[Me2]1[Na] | 5[Cys]1[Ox] |
| 610.44 | 2[Cys]3[Ph-OH]2[Me2]1[Na] | 1[Cys]5[Ph-OH] |
| 613.79 | 3[Ph-OH]4[Ph] | 2[Cys]3[Ph-OH]1[Ph] |
| 613.79 | 5[Ph-OH]1[Ph]2[Na] | 3[Ph-OH]4[Ph] |
| 613.79 | 1[Ph-OH]6[Ph]1[Na]1[Me1] | 4[Cys]1[Ph]1[Me3]1[Me1] |
| 633.73 | 5[Cys]1[Na]1[Ox] | 3[Cys]2[Ph-OH]1[Ph] |
| 633.73 | 3[Cys]1[Ph]3[Me3]1[Ac]1[Me2] | 5[Cys]1[Na]1[Ox] |
| 633.73 | 3[Cys]1[Ph]2[Me3]2[Ac]1[Me2] | 2[Cys]3[Ph-OH]1[Ph]1[Na] |
| 652.47 | 4[Cys]2[Ph]1[Ox] | 3[Cys]3[Ph-OH] |
| 652.47 | 2[Cys]3[Ph-OH]1[Me3]2[Me2]1[Na] | 4[Cys]2[Ph]1[Ox] |
| 652.47 | 2[Cys]3[Ph-OH]2[Me3]1[Na]1[Me1] | 5[Cys]1[Me3]1[Ox] |
| 692.56 | 4[Cys]2[Ph]1[Ac]1[Me1] | 5[Cys]1[Ph-OH] |
| 692.56 | 4[Cys]1[Ph-OH]1[Ph]1[Na]1[Ox] | 2[Cys]3[Ph-OH]2[Ph] |
| 692.56 | 4[Cys]2[Ph]1[Me3]1[Me1] | 3[Cys]3[Ph-OH]1[Ac] |
| 711.78 | 5[Cys]1[Ph]1[Na]1[Me1] | 2[Cys]4[Ph-OH]1[Ph] |
| 711.78 | 5[Cys]1[Me3]2[Na]1[Ox]1[Me1] | 5[Cys]1[Ph]1[Na]1[Me1] |
| 711.78 | 4[Ph-OH]4[Ph] | 3[Cys]1[Ph-OH]3[Ph]1[Ox] |
| 731.75 | 5[Cys]1[Ph]1[Ac]1[Me1] | 3[Cys]3[Ph-OH]1[Ph] |
| 731.75 | 5[Cys]1[Ph]1[Me3]1[Me1] | 5[Cys]1[Ph]1[Ac]1[Me1] |
| 731.75 | 5[Cys]1[Ph]2[Me2] | 5[Cys]1[Ph]1[Me3]1[Me1] |
| 792.89 | 4[Cys]1[Ph-OH]3[Me3]1[Ac]1[Me2]1[Na] | 5[Cys]2[Ph-OH] |
| 792.89 | 4[Cys]1[Ph-OH]4[Me3]1[Me2]1[Na] | 4[Cys]3[Ph-OH]1[Na] |
| 792.89 | 4[Cys]1[Ph-OH]2[Me3]2[Ac]1[Me2]1[Na] | 3[Cys]2[Ph-OH]3[Ph] |
| 812.52 | 4[Cys]3[Ph-OH]1[Ac] | 4[Cys]3[Ph-OH]1[Ac] |
| 812.52 | 4[Cys]3[Ph-OH]1[Me3] | 4[Cys]3[Ph-OH]1[Me3] |
| 812.52 | 2[Cys]5[Ph-OH]2[Me3] | 3[Cys]3[Ph-OH]2[Ph] |
